# A population-level statistic for assessing Mendelian behavior of genotyping-by-sequencing data from highly duplicated genomes

**DOI:** 10.1101/2020.01.11.902890

**Authors:** Lindsay V. Clark, Wittney Mays, Alexander E. Lipka, Erik J. Sacks

## Abstract

**Background:** Given the economic and environmental importance of allopolyploids and other species with highly duplicated genomes, there is a need for methods to distinguish paralogs, i.e. duplicate sequences within a genome, from Mendelian loci, i.e. single copy sequences that pair at meiosis. The ratio of observed to expected heterozygosity is an effective tool for filtering loci but requires genotyping to be performed first at a high computational cost, whereas counting the number of sequence tags detected per genotype is computationally quick but very ineffective in inbred or polyploid populations. Therefore, new methods are needed for filtering paralogs.

**Results:** We introduce a novel statistic, *H*_*ind*_/*H*_*E*_, that uses the probability that two reads sampled from a genotype will belong to different alleles, instead of observed heterozygosity. The expected value of *H*_*ind*_/*H*_*E*_ is the same across all loci in a dataset, regardless of read depth or allele frequency. In contrast to methods based on observed heterozygosity, it can be estimated and used for filtering loci prior to genotype calling. In addition to filtering paralogs, it can be used to filter loci with null alleles or high overdispersion, and identify individuals with unexpected ploidy and hybrid status. We demonstrate that the statistic is useful at read depths as low as five to 10, well below the depth needed for accurate genotype calling in polyploid and outcrossing species.

**Conclusions:** Our methodology for estimating *H*_*ind*_/*H*_*E*_ across loci and individuals, as well as determining reasonable thresholds for filtering loci, is implemented in polyRAD v1.6, available at https://github.com/lvclark/polyRAD. In large sequencing datasets, we anticipate that the ability to filter markers and identify problematic individuals prior to genotype calling will save researchers considerable computational time.

## Background

Highly duplicated genome sequences are common throughout the plant kingdom. These include recent allopolyploids such as wheat, cotton, canola, strawberry, and coffee, as well as species with evidence of ancient whole genome duplication such as maize and legumes [1]. This phenomenon is also present in the animal kingdom, for example allopolyploidy in the model frog *Xenopus*, as well as an ancient tetraploidization event followed by diploidization in salmonid fishes [2, 3]. For species in which paralogous sequences no longer pair at meiosis, accurate separation of paralogs in DNA and RNA sequence analysis, including reference genome assembly, remains challenging [4]. This separation of paralogs is especially important in variant calling, because SNPs and indels will not behave in a Mendelian fashion if the reads originate from more than one locus yet are erroneously attributed to a single locus [5]. Accurate variant calling therefore impacts all downstream analysis that assumes Mendelian inheritance, including linkage and QTL mapping, association studies, genomic selection, population genetics, and parentage analysis. For example, failure to remove paralogs from downstream analysis has been demonstrated to bias estimates of allele frequency and inbreeding as well as population structure [4, 6–8].

Due in part to the difficulty of assembling highly duplicated reference genomes, several methods have been published for filtering collapsed paralogous loci from genotyping-by-sequencing (GBS, including restriction-site associated DNA sequencing (RAD) approaches) datasets without the need for a reference genome. The most straightforward approach is to call genotypes and then determine if observed heterozygosity exceeds expected heterozygosity [9–11]. However, sampling error at low read depth can confound this filtering step by causing heterozygotes to be miscalled as homozygotes, lowering the observed heterozygosity. Moreover, estimating observed heterozygosity becomes complicated when polysomic inheritance is expected, due to the challenge of estimating allele copy number. Bayesian genotype calling methods mitigate the underestimation of observed heterozygosity, but at substantial computational cost [12–14]. Another approach is to filter loci that have read depth above an arbitrary threshold [15], although due to differences in amplification efficiency based on fragment size and GC content, this method could fail to filter some paralogs while filtering other non-paralogous loci. Peterson et al. [16] developed a method, extended by Willis et al. [17], that involved counting the number of unique haplotypes per individual for a putative locus, with the idea that in a collapsed paralog, the number of haplotypes would exceed the ploidy. However, this method can be confounded by sequencing error and cross-contamination among samples, and its sensitivity depends on allele frequencies, inbreeding, and ploidy. Other approaches have examined read depth ratios within individual genotypes [18] as well as read depth ratios in combination with observed heterozygosity [19]. Lastly, multiple methods identify putative paralogs based on networks of similarity among sequence tags [6, 20].

We present a novel statistic, *H*_*ind*_/*H*_*E*_, for evaluating marker quality, in particular for assessing whether a marker represents one Mendelian locus or multiple collapsed paralogous loci, based upon read depth distribution in a population. For a Mendelian locus, the statistic has the same expected value regardless of number of alleles, allele frequency, and total read depth. As a result, the distribution of the statistic can be visualized across loci in order to identify threshold values for filtering. The expected value can be calculated from ploidy (assuming disomic or polysomic inheritance) and the inbreeding coefficient, or the mode value of the statistic in a population can be used to estimate ploidy or inbreeding. Notably, because genotype calls are not needed in order to estimate this statistic, it can be used for filtering loci before any genotype calling is performed, saving computation time. Technical parameters such as sequencing error and overdispersion can influence estimates, but are explored here using simulated data so that they can be accounted for. We extend our Bayesian genotype calling software polyRAD [12] to implement the novel statistic and determine appropriate cutoffs.

## Results

### The *H*_*ind*_ statistic

Here we describe a novel statistic, *H*_*ind*_, that is based on sequence read depth across all alleles at a given locus and sample, and is agnostic of genotype calls, inheritance mode, and ploidy. It is related to observed heterozygosity, *H*_*O*_, which in a diploid can be thought of as a matrix of ones and zeros indicating whether the genotype at each sample*locus is heterozygous. *H*_*ind*_ is instead a number ranging from zero to one, indicating the probability that if two sequencing reads were sampled without replacement at that sample*locus, they would represent different alleles. The abbreviation “ind” stands for “individual”, as it is calculated for each individual before averaging across a population. It can be calculated for SNP loci or for multiallelic haplotype- or tag-based loci, as long as allelic read depth is available.

The expected value for *H*_*ind*_ in a natural population of diploids or polysomic polyploids is:

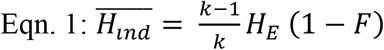

where *k* is the ploidy, *H*_*E*_ is the expected heterozygosity at the same locus, and *F* is the inbreeding coefficient. *H*_*E*_ is the probability that two alleles drawn at random from the population will be different, (1 – *F*) is the probability that two alleles randomly drawn from an individual will not be identical by descent, and (*k* – 1)/*k* is the probability that two sequencing reads originated from different chromosome copies. Multiplied together, these three terms yield the probability that two sequence reads from one sample at one locus will be different from each other.

If we divide *H*_*ind*_ by *H*_*E*_:

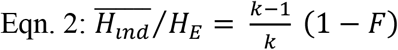

we now have a statistic that is only dependent on ploidy and inbreeding, two parameters that we will assume to be consistent across samples and loci.

In a mapping population, the term *H*_*E*_ * (1 – *F*) must be replaced by the probability, for a given locus, that two locus copies in a progeny will be different alleles. This requires knowledge of the ploidy, parental genotypes, and population design including number of generations of backcrossing and self-fertilization. This probability, which we will call *H*_*E*.*map*_, can be estimated by simulation of the cross. The expectation is then:

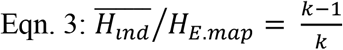

Common factors that influence *H*_*ind*_/*H*_*E*_ are listed in Table 1, and explored in subsequent sections.

**Table 1.** Biological and technical parameters that influence the expected value and variance of *H*_*ind*_/*H*_*E*_.

| Parameter | Effect |
| --- | --- |
| Ploidy | Expected value increases with ploidy. |
| Inbreeding (including population structure) | Expected value decreases as inbreeding increases. |
| Hybridization | Value increases with increase in heterozygosity from hybridization across species or divergent populations. |
| Paralogy | Value increases if multiple loci are collapsed into one. |
| Overdispersion | Expected value decreases as read depth ratios deviate further from allelic dosage. |
| Sequencing error | Value is biased upward by sequencing error, especially at low minor allele frequencies. |
| Null alleles (e.g. restriction site polymorphisms, deletions) | Expected value decreases with increasing null allele frequency. |
| Minor allele frequency | Variance decreases at increased minor allele frequency. Overdispersion, sequencing error, or very low read depth in combination with low minor allele frequency bias the value upward. |
| Sample size | Variance decreases at increased sample size. |
| Number of alleles | Multiallelic loci have lower variance than biallelic SNPs. |
| Read depth | Low read depth loci tend to have low values due to the presence of null alleles. High read depth loci tend to have high values due to paralogy. Genome-wide increases in read depth (e.g. larger library size) reduce variance in the statistic, as well as reducing upward bias at low minor allele frequencies. |
| Polyploid mapping populations | Variance is lower at markers with higher heterozygosity in the progeny. |

### Empirical estimation of *H*_*ind*_/*H*_*E*_

Say that we have sequence read depths, {*d*_1*m*_ … *d*_*jm*_}, across a set of *j* alleles at a single locus in an individual *m*. Total read depth in one individual is

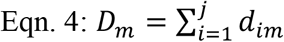

As long as there are two or more reads, we can estimate *H*_*ind*_ within that individual using the Gini-Simpson index [21]:

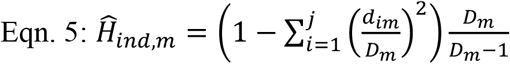

For a population of *n* individuals with sequencing reads, allele frequencies are estimated from average within-individual read depth ratios:

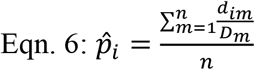

And expected heterozygosity is estimated as

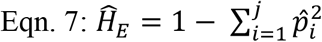

Averaged across *n* individuals with two or more reads at a given locus in a natural population, the expectation is then:

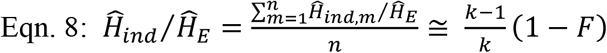

In a mapping population, 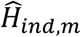 is estimated in the same way. *H*_*E*.*map*_ is estimated from parental genotypes and population design, and the expected average ratio within a locus is given in Eqn. 3.

### Utility of *H*_*ind*_/*H*_*E*_ for detecting collapsed paralogs in a diversity panel

To compare the distribution of *H*_*ind*_/*H*_*E*_ values for Mendelian loci versus collapsed paralogs, we aligned *M. sacchariflorus* tag sequences to the *M. sinensis* reference genome, in which they should align to the correct paralog most of the time, and to the *S. bicolor* reference genome, in which two paralogs from *Miscanthus* correspond to one alignment location. We found that loci with a mean read depth less than five had very low estimates of *H*_*ind*_/*H*_*E*_, likely due to restriction site polymorphisms or other technical issues (Fig. 1 and Additional File 1: Fig. S1). As mean read depth increased above 100 in our dataset, however, loci tended to have *H*_*ind*_/*H*_*E*_ values above the expectation for a Mendelian locus, suggesting that most loci at this depth and higher were in fact collapsed paralogs (Fig. 1, and Additional File 1: Figs. S1 and S2).

**Figure 1.**
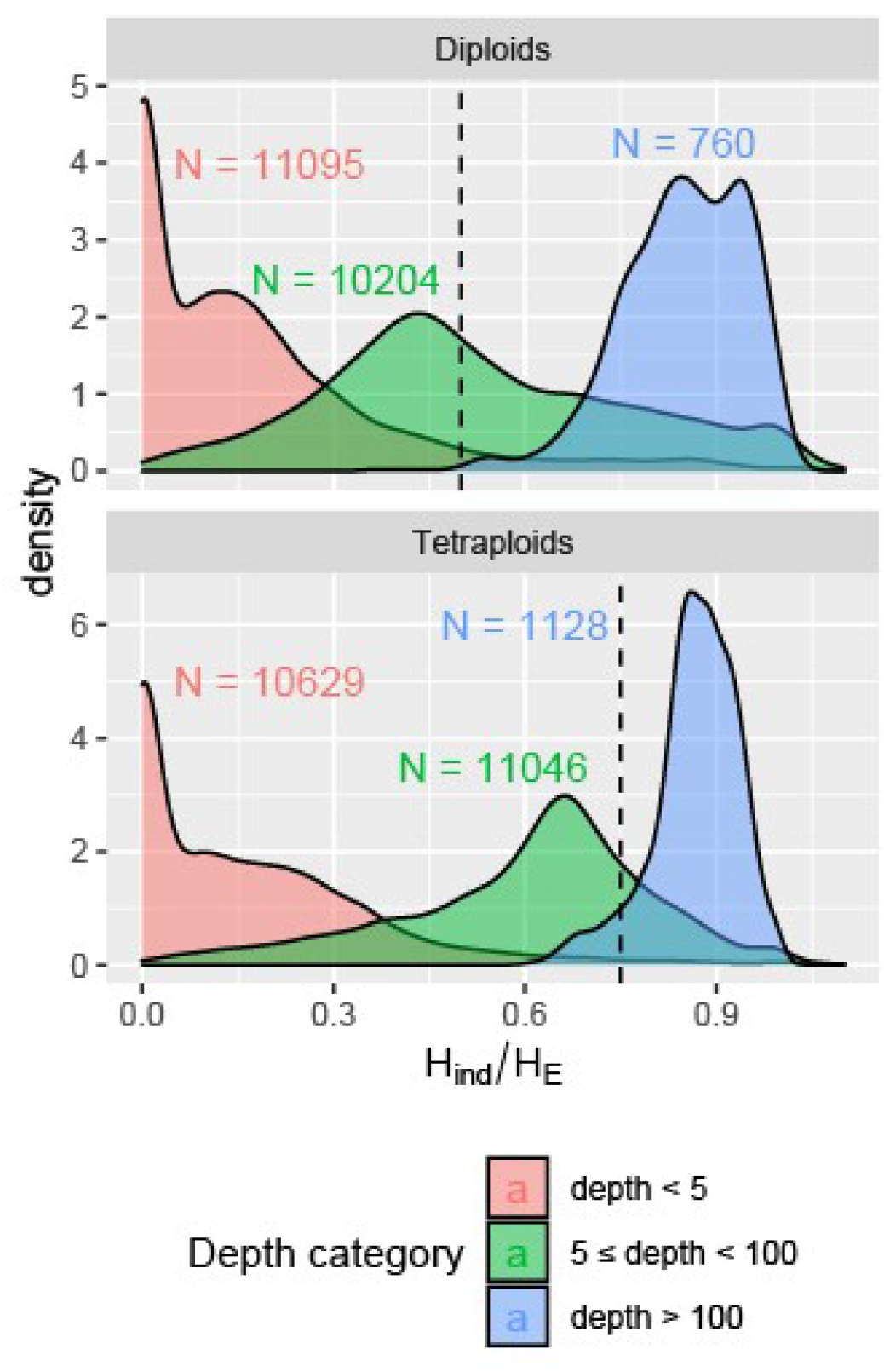
Relationship between *H*_*ind*_/*H*_*E*_ statistic and mean sequence read depth per locus. Loci were called across 356 diploid and 268 tetraploid *Miscanthus sacchariflorus* based on alignments to the *M. sinensis* reference genome. The number of loci in each depth category is indicated. Fig. S1 in Additional File 1 provides justification for the depth thresholds for categories. The expected value for a Mendelian locus in Hardy-Weinberg equilibrium is shown with a dashed line.

When a mean depth of five was used as a cutoff and the *M. sinensis* genome was used as a reference, the peak *H*_*ind*_/*H*_*E*_ value was slightly below the expected values of 0.5 for diploids and 0.75 for tetraploids (Fig. 2), indicating some inbreeding, likely due to population structure [22]. A second peak was observed at a higher value of *H*_*ind*_/*H*_*E*_ (Fig. 2), likely representing sets of tags that belonged to different Mendelian loci despite aligning to the same location (i.e. misalignments). When *S. bicolor* was used as the reference genome, the opposite trend was observed, where most loci had a *H*_*ind*_/*H*_*E*_ above the expected value, indicating collapsed paralogs, but a second peak was observed closer to the expected value, indicating regions in the *S. bicolor* genome that may only have synteny with one region of the *M. sinensis* genome (Fig. 2).

**Figure 2.**
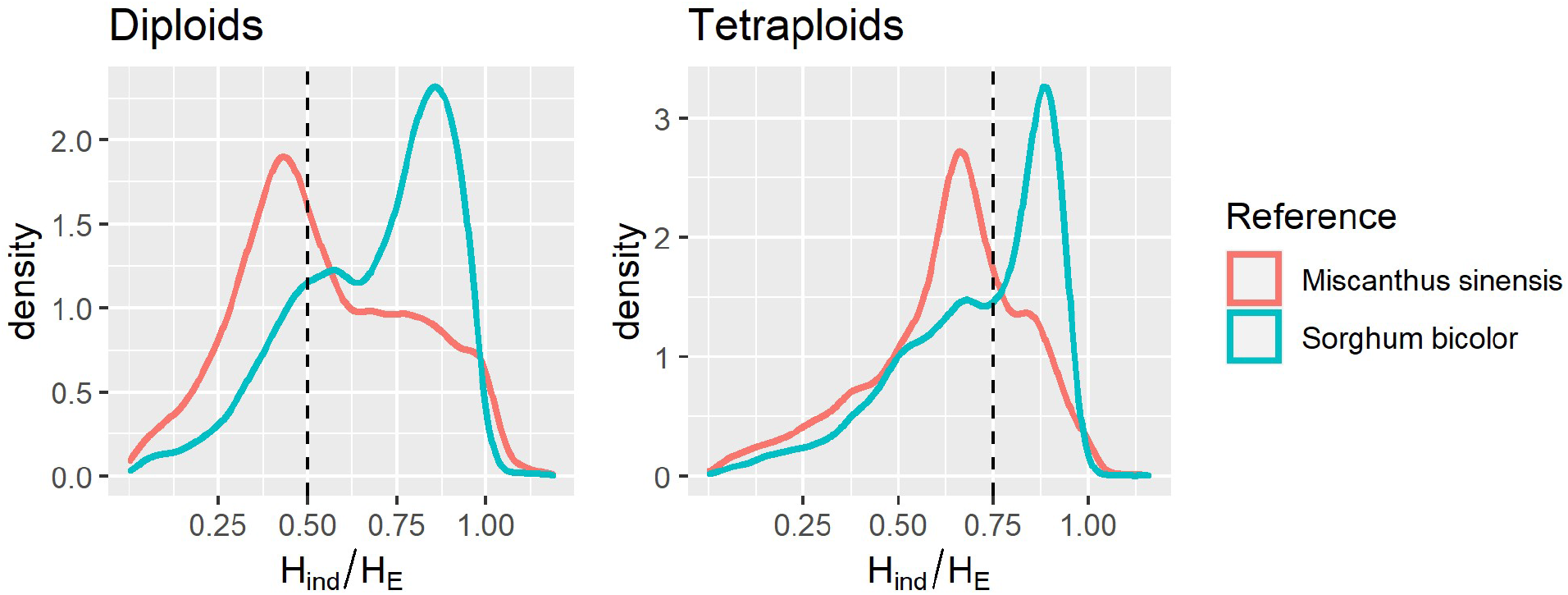
Effect of reference genome and ploidy on *H*_*ind*_/*H*_*E*_ per locus in *Miscanthus sacchariflorus*. Loci with a mean read depth below five were omitted, leaving 11,516 loci aligned to the *M. sinensis* reference and 8,820 loci aligned to the *Sorghum bicolor* reference. Expected values for Mendelian loci under Hardy-Weinberg equilibrium are shown with dashed lines.

Although the peaks overlapped somewhat, they were distinct enough that a reasonable threshold for identifying putative collapsed paralogs could be visually determined (Fig. 2). Moreover, although the diploid and tetraploid datasets were processed separately, they were largely in agreement about which loci were Mendelian and which were collapsed paralogs (Additional File 1: Fig. S2), suggesting that the filtering performed in one population can be applied to another population, which could be especially useful for populations that are too small for accurate estimation of *H*_*ind*_/*H*_*E*_.

In both the diploid and tetraploid datasets, the distribution and peak values of *H*_*ind*_/*H*_*E*_ were similar regardless of whether biallelic SNPs or multiallelic, haplotype-based markers were used (Additional File 1: Fig. S3). However, the variance of *H*_*ind*_/*H*_*E*_ was approximately 20% higher when SNPs were used, suggesting that the higher information content of multiallelic markers improves the precision of *H*_*ind*_/*H*_*E*_ estimates.

The *ExpectedHindHe* function in polyRAD was used to set thresholds for filtering the diploid and tetraploid datasets. Based on results from the *TestOverdispersion* function, the overdispersion parameter was set to 11 for diploids and 10 for tetraploids. Based on the observed distribution of *H*_*ind*_/*H*_*E*_ in the dataset, the inbreeding coefficient was set to 0.35 for diploids and 0.25 for tetraploids. Based on these parameters, as well as read depth and allele frequencies in the datasets, the ranges for retaining 95% of Mendelian loci were 0.175 to 0.584 in diploids and 0.356 to 0.716 in tetraploids as estimated by *ExpectedHindHe*, resulting in 40.2% and 42.3% of loci being filtered, respectively (Table 2). Markers within genes were underrepresented among markers that were filtered for having *H*_*ind*_/*H*_*E*_ below the lower threshold, and overrepresented among markers that were filtered for having *H*_*ind*_/*H*_*E*_ above the upper threshold, significant in Fisher’s Exact Test at P < 0.0005 (Table 2). Markers that were filtered having *H*_*ind*_/*H*_*E*_ above the upper threshold tended to have minor allele frequencies that were very low, consistent with the markers representing sequencing error rather than true alleles, or very high, consistent with the markers representing collapsed paralogs (Fig. 3).

**Table 2.** Contingency tables of number of markers retained and filtered for being above or below *H*_*ind*_/*H*_*E*_ thresholds in *Miscanthus sacchariflorus*, by whether or not the marker was within a gene.

|  | Diploids |  | Tetraploids |  |
| --- | --- | --- | --- | --- |
|  | In a gene | Not in a gene | In a gene | Not in a gene |
| Filtered; too low | 337 | 950 | 588 | 1727 |
| Retained | 2201 | 3654 | 2419 | 3500 |
| Filtered; too high | 1361 | 1287 | 1091 | 930 |

**Figure 3.**
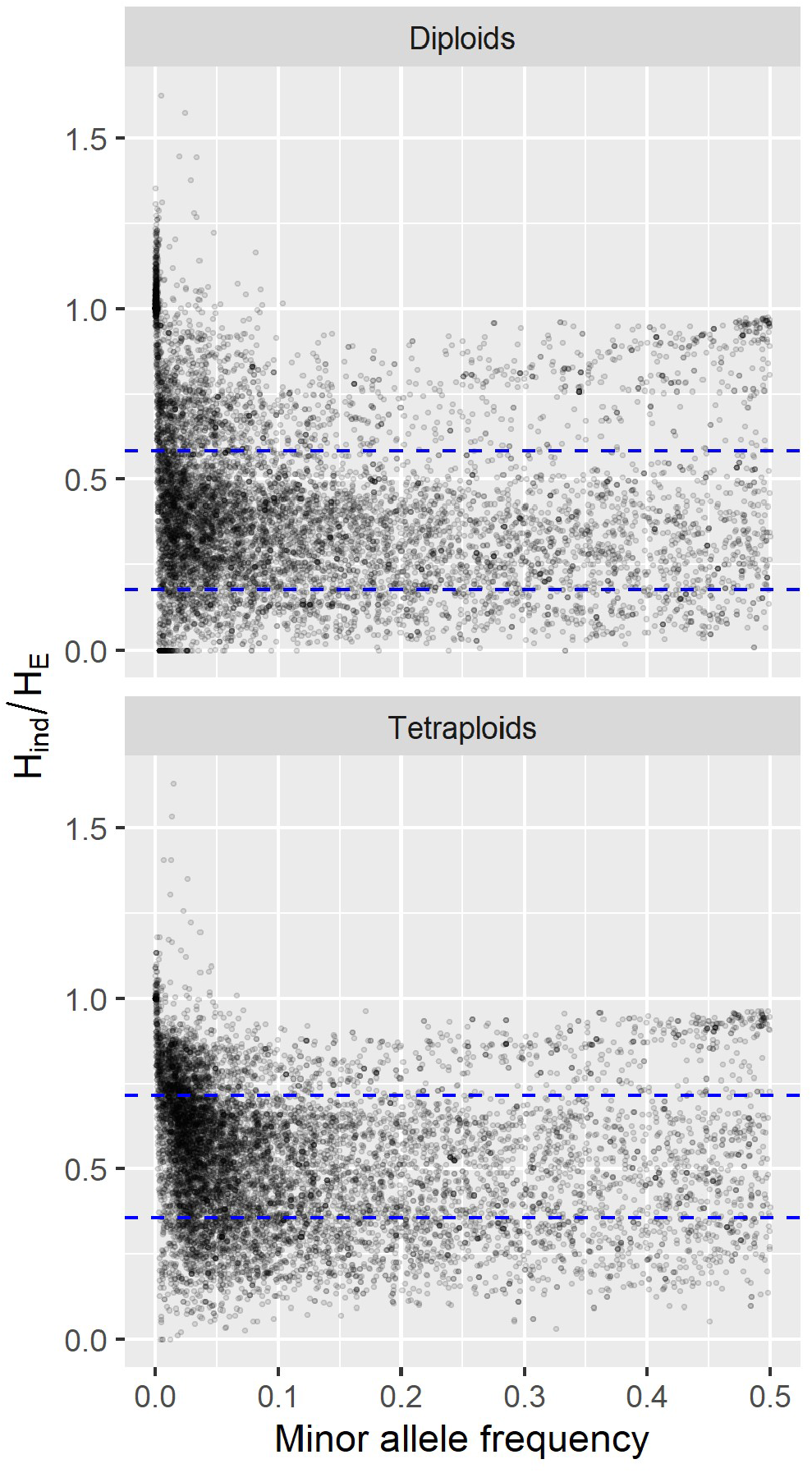
Filtering by *H*_*ind*_/*H*_*E*_ vs. minor allele frequency in *Miscanthus sacchariflorus*. A dataset of 10,458 SNP loci was tested across 356 diploid and 268 tetraploid individuals. Blue dashed lines indicate filtering thresholds to retain 95% of Mendelian loci based on simulated distributions.

### By individual, *H*_*ind*_/*H*_*E*_ reflects ploidy and hybrid status

In addition to evaluating the mean *H*_*ind*_/*H*_*E*_ within loci, we also obtained the mean statistic within individuals in order to assess the utility of the statistic for determining ploidy. We found that *H*_*ind*_/*H*_*E*_ increased with ploidy, largely independent of read depth (Fig. 4). Although the distributions overlapped too much for *H*_*ind*_/*H*_*E*_ to be a conclusive indicator of ploidy, it could still potentially be used to identify outlier individuals whose ploidy should be confirmed by other means (e.g. flow cytometry). Additionally, because our empirical dataset included many natural interspecific (*M. sacchariflorus* × *M. sinensis*) F1 hybrid and backcross individuals, we were also able to observe that *H*_*ind*_/*H*_*E*_ values were considerably higher in hybrids than in non-hybrids, reflecting higher heterozygosity (Fig. 4).

**Figure 4.**
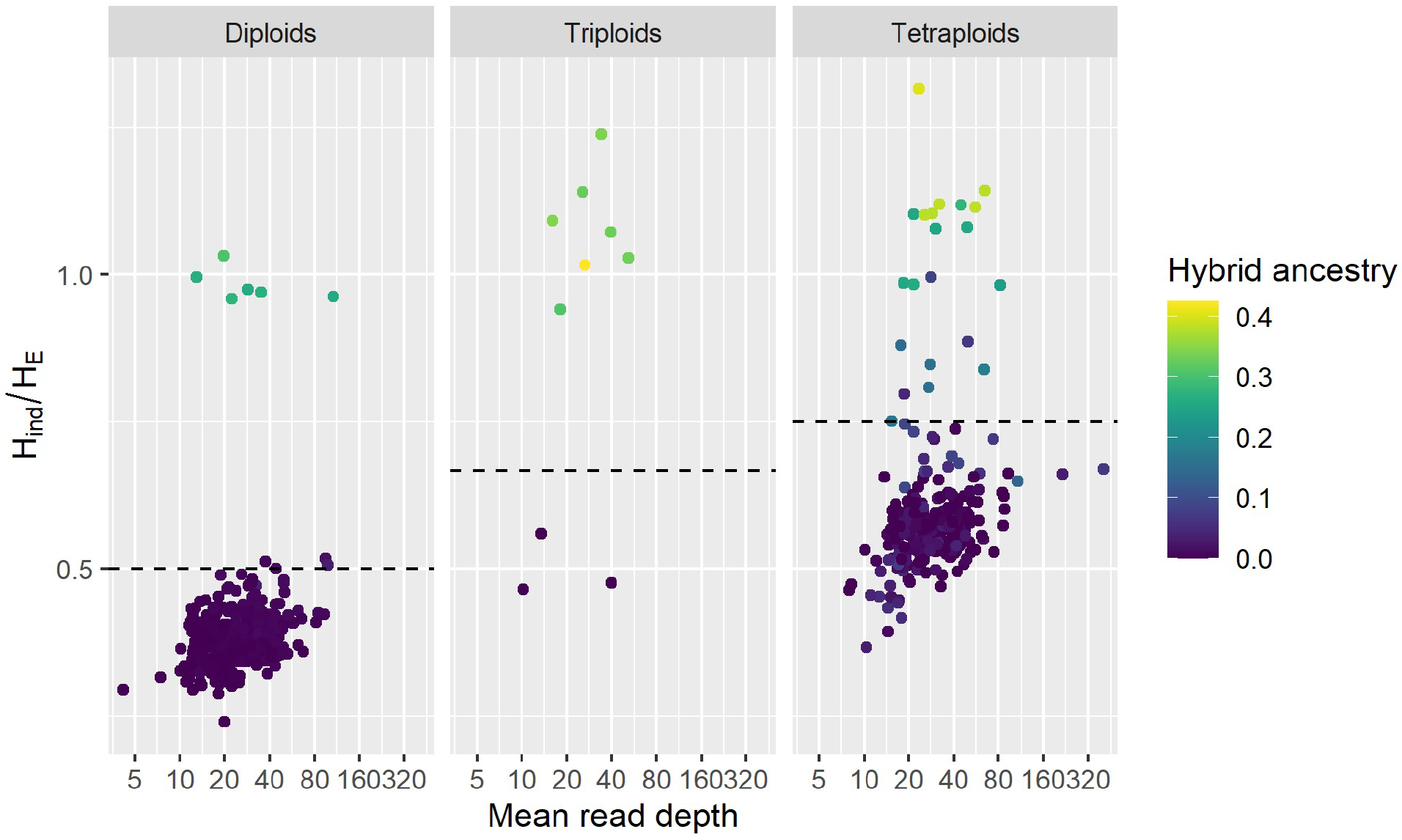
Relationship between ploidy, sequence read depth, hybrid ancestry, and *H*_*ind*_/*H*_*E*_ among 620 *M. sacchariflorus* individuals. Ploidy and proportion of ancestry from *M. sinensis* (hybrid ancestry) were determined previously [22]. Read depth and *H*_*ind*_/*H*_*E*_ were averaged across 10,000 loci. The expected value for *H*_*ind*_/*H*_*E*_ under Hardy-Weinberg equilibrium is shown with the dashed line.

### Variance and bias in the *H*_*ind*_/*H*_*E*_ statistic using simulated data

Using simulated data resembling a diversity panel or natural population, the mean *H*_*ind*_/*H*_*E*_ estimate decreased as inbreeding increased, with diploid and tetraploid loci being indistinguishable at an inbreeding coefficient of 0.8 or higher (Fig. 5). Sequencing error had little effect on the estimate at a minor allele frequency of 0.05, but caused an inflated estimate at a minor allele frequency of 0.01, particularly as inbreeding increased (Fig. 5). Variance and bias in the statistic were minimized if there were at least 500 samples, minor allele frequency was 0.05 or higher, and read depth was at least 5 (Fig. 6). Ploidy had negligible impact on variance and bias (Fig. 6). Read depth and minor allele frequency influenced the estimates for collapsed paralogs, but not enough to interfere with distinguishing them from Mendelian markers (Fig. 6). As expected, overdispersion (deviation of read depth ratios from allelic dosage ratios) reduced the mean *H*_*ind*_/*H*_*E*_ estimate, with the effect of overdispersion being greater at higher minor allele frequencies (Additional File 1: Fig. S4). The *H*_*ind*_/*H*_*E*_ estimate also decreased linearly as null allele frequency increased (Additional File 1: Fig. S5).

**Figure 5.**
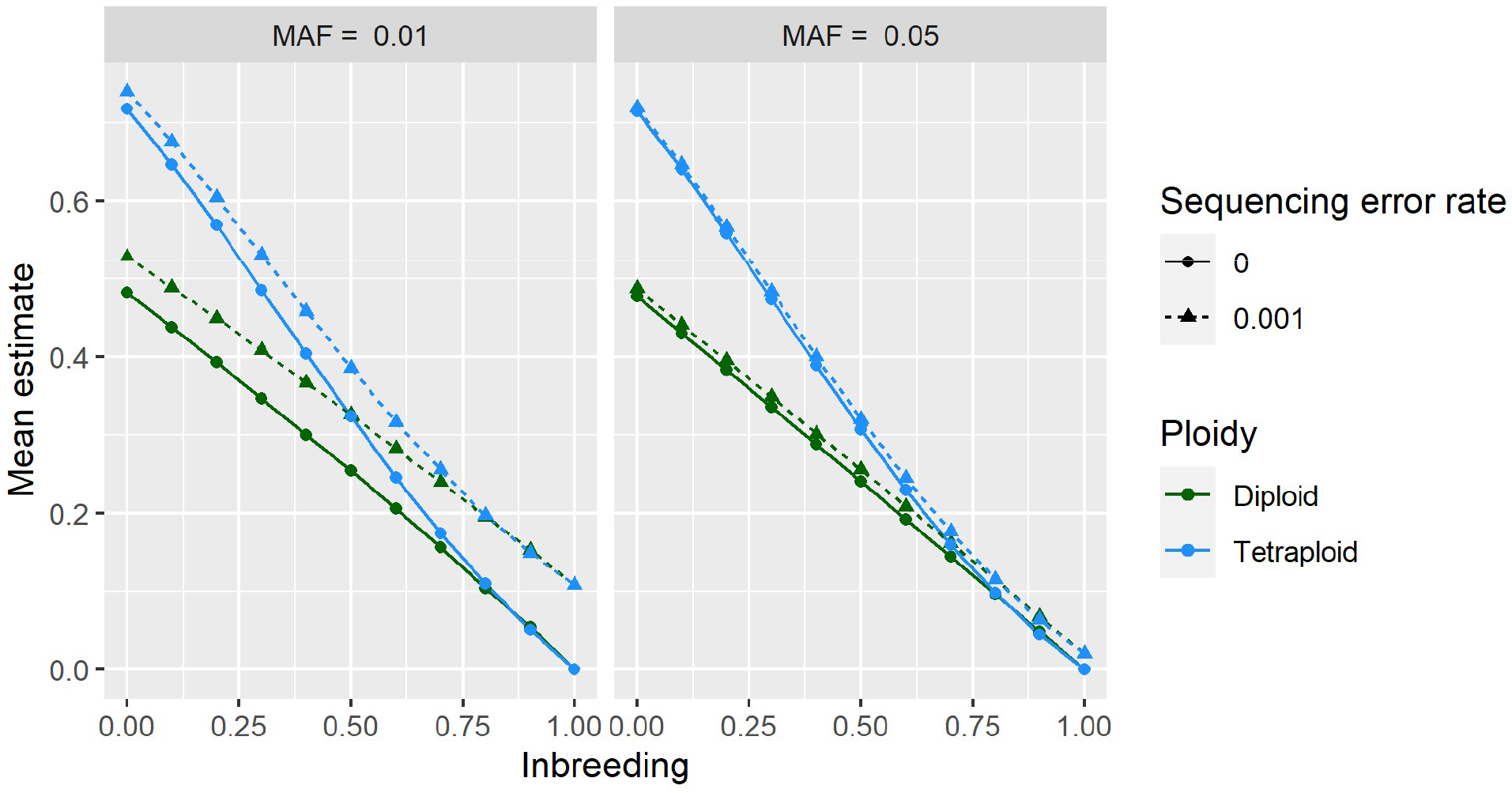
Combined effects of inbreeding, ploidy, minor allele frequency (MAF), and sequencing error on mean estimates of *H*_*ind*_/*H*_*E*_ using simulated data. At each combination of parameters, 20,000 biallelic loci were simulated with a read depth of 20 and overdispersion parameter of 20. The x-axis indicates the inbreeding coefficient (the probability that two alleles in an individual are identical by descent) while the y-axis indicates the *H*_*ind*_/*H*_*E*_ estimate.

**Figure 6.**
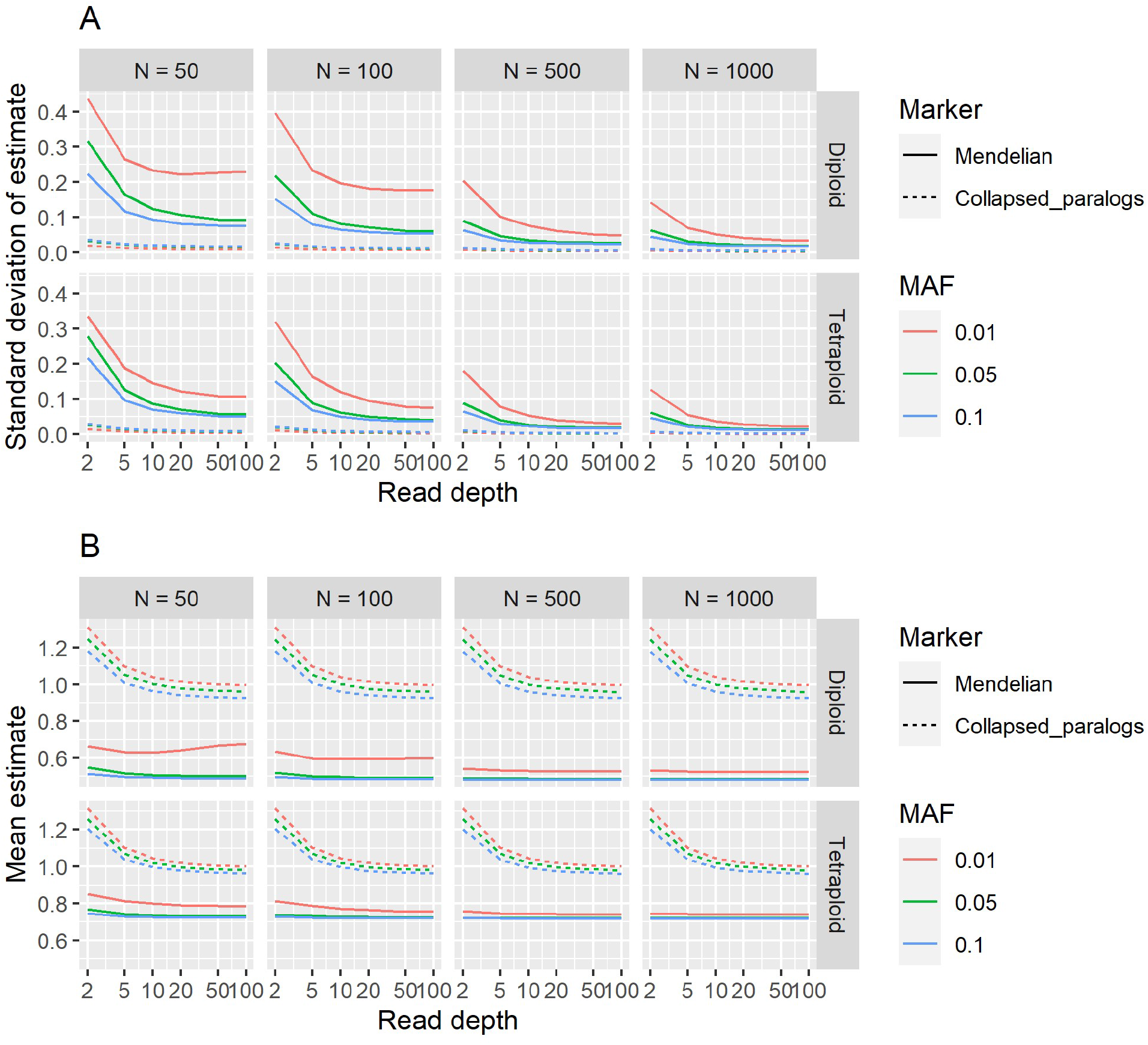
Effect of sample size, read depth, and minor allele frequency on variance and bias of estimates of *H*_*ind*_/*H*_*E*_. For each combination of ploidy, sample size (N), read depth, and minor allele frequency (MAF), 5000 biallelic Mendelian loci were simulated under Hardy-Weinberg Equilibrium with an overdispersion parameter of 20 and sequencing error rate of 0.001. Additionally, 5000 collapsed paralogs, each consisting of two Mendelian loci, were simulated under each set of the same parameters. (A) Standard deviation of *H*_*ind*_/*H*_*E*_ estimates. (B) Mean *H*_*ind*_/*H*_*E*_ estimates. Expected values are 0.5 for diploids and 0.75 for tetraploids; deviations from these values indicate bias in estimation.

In simulated F1 mapping populations, the standard deviation of the *H*_*ind*_/*H*_*E*_ ranged from 0.012 to 0.076 depending on the marker type (Fig. 7). In tetraploids, marker types with high expected heterozygosity in the progeny, such as triplex x nulliplex and triplex x simplex, had lower variance in the estimate than marker types with lower expected heterozygosity in the progeny, such as simplex x nulliplex and simplex x simplex (Fig. 7). A few rare markers had *H*_*ind*_/*H*_*E*_ estimates that deviated very far from the expected value, indicating that the parents were incorrectly genotyped (Fig. 7).

**Figure 7.**
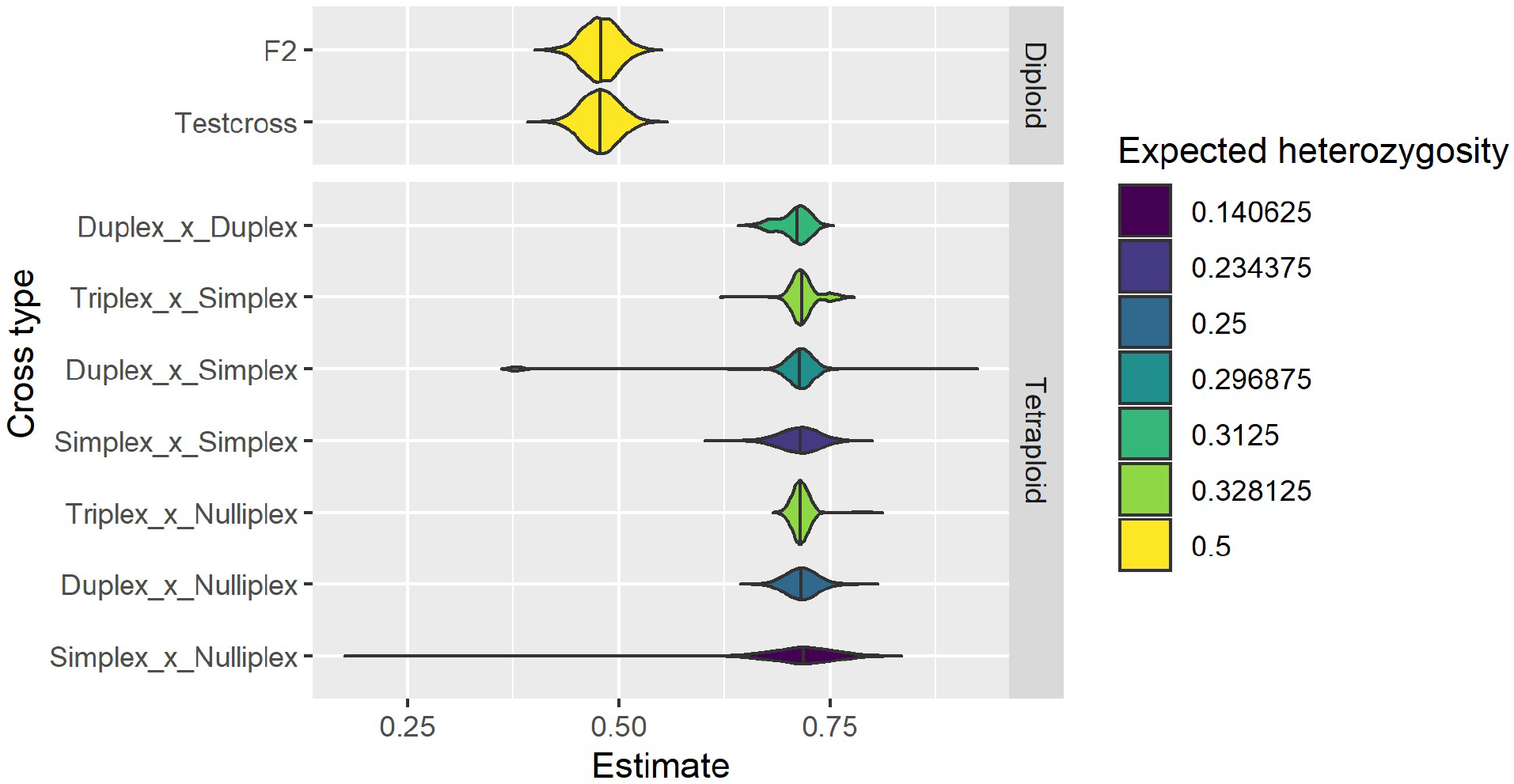
Distribution of *H*_*ind*_/*H*_*E*_ estimates in simulated F1 mapping populations. For each cross type, 5000 biallelic loci with a read depth of 20, overdispersion parameter of 20, and sequencing error rate of 0.001 were simulated across 500 individuals.

### Comparison with other approaches

To compare effectiveness at filtering paralogs between *H*_*ind*_/*H*_*E*_ and other approaches, 1000 Mendelian loci and 1000 collapsed paralogs were simulated in 200 diploid and 200 tetraploid individuals at three levels of inbreeding. The median allele frequency was 0.026 and median read depth per Mendelian locus was 21. For each statistic, the 95^th^ percentile for Mendelian loci was determined, and the proportion of collapsed paralogs that would be filtered at that threshold was estimated. The *H*_*ind*_/*H*_*E*_ approach and observed over expected heterozygosity (*H*_*O*_/*H*_*E*_) performed best, with *H*_*O*_/*H*_*E*_ having the disadvantage that genotyping must be performed before it can be estimated, thus increasing processing time two orders of magnitude over *H*_*ind*_/*H*_*E*_ (Table 3). The *H*_*ind*_/*H*_*E*_ thresholds used for filtering were 0.58, 0.41, and 0.17 in diploids and 0.76, 0.48, and 0.17 in tetraploids at inbreeding levels of 0.1, 0.5, and 0.9, respectively. The haplotype counting approach [17] and allelic depth ratio Z-score approach [19] both performed reasonably well in diploids but were much less effective in tetraploids, with haplotype counting being useless in tetraploids at high inbreeding, while the Z-score approach additionally suffered in terms of computational time due to the need for genotyping. However, haplotype counting used 2- to 3-fold less computational time than *H*_*ind*_/*H*_*E*_, and thus could be advantageous in diploids when millions of loci are being processed. Lastly, filtering on read depth alone was not very effective given the variation in read depth among loci.

**Table 3.**
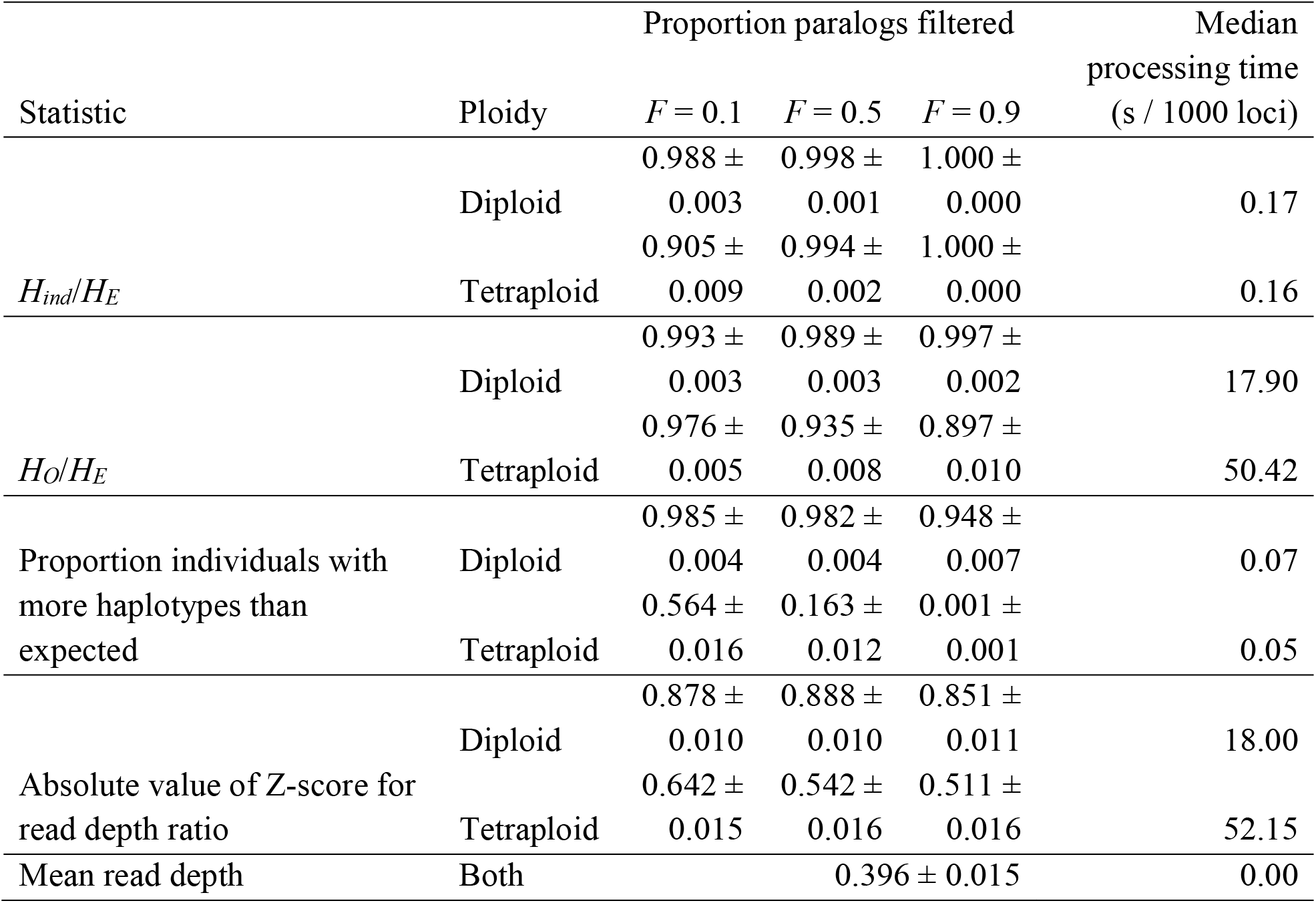
Effectiveness of various statistics for identifying paralogs, using simulated data across three levels of inbreeding. Standard error is shown for proportion paralogs filtered.

## Discussion

### Properties of the *H*_*ind*_/*H*_*E*_ statistic

While *H*_*ind*_/*H*_*E*_ can be used, in combination with other metrics, to assess locus quality, this should be performed with an understanding of what biological and technical phenomena can cause it to deviate from the expected value. Inbreeding from any source will lower the expected value below (*k* – 1)/*k*, where *k* is the ploidy; this includes not only self-fertilization and preferential mating with relatives, but also population structure, which is why we observed values below (*k* – 1)/*k* even in self-incompatible, wind-pollinated *M. sacchariflorus* (Figs. 1-4). A benefit of this, however, is that as long as ploidy is known and overdispersion can be reasonably estimated (e.g. with the *TestOverdispersion* function in polyRAD), *H*_*ind*_/*H*_*E*_ can be used to estimate inbreeding, either at the population or individual level, directly from sequence read depth. Given that we observed *H*_*ind*_/*H*_*E*_ to be inflated at low minor allele frequencies, we recommend using the mode *H*_*ind*_/*H*_*E*_ at markers with minor allele frequency of at least 0.05 for estimating inbreeding. Additionally, individuals that are hybrids between species or between highly diverged populations, as well as DNA samples that are an accidental mix of two or more individuals, may have *H*_*ind*_/*H*_*E*_ above the expected value (Fig. 4). Strong selection for homozygotes or heterozygotes at particular loci would be expected to lower and raise *H*_*ind*_/*H*_*E*_, respectively.

At the locus level, a *H*_*ind*_/*H*_*E*_ that exceeds the expected value can be an indication that alleles are derived from paralogous loci rather than a true Mendelian locus. More broadly, if all alleles truly belong to a single locus, then the expected value is (1 – *F*)(*k* – 1)/*k*. However, if a set of random, independent alleles were assigned to one putative locus, the expected value of *H*_*ind*_/*H*_*E*_ would be one, because the probability of sampling reads from two different alleles within one individual would be the same as the probability of sampling reads from two different alleles in the general population. In the *M. sacchariflorus* dataset, markers within genes were overrepresented among markers that were filtered for having *H*_*ind*_/*H*_*E*_ above the expected value, likely due to high sequence conservation between paralogs (Table 2). A *H*_*ind*_/*H*_*E*_ of zero could indicate a cytoplasmic marker, because while there may be variation in the population, each individual would only be expected to possess reads from one allele. Loci with highly overdispersed read depth distributions due to technical issues such as differential fragment size or variation in library preparation would also be expected to have *H*_*ind*_/*H*_*E*_ below expectations; it may be advantageous to filter these from the dataset as they will tend to yield poor-quality genotype calls. Lastly, loci with common null alleles have lower than expected *H*_*ind*_/*H*_*E*_ (Additional File 1: Fig. S3), resulting in a tendency to filter loci that are not within genes as these regions are less conserved (Table 2). Null alleles can be the result of restriction cut site polymorphism in RAD-based techniques, primer binding site mutations in amplicon sequencing, or deletion mutations using any genotyping method. Because they are a common problem, *H*_*ind*_/*H*_*E*_ can be used to identify and filter loci with null alleles.

The expected value of *H*_*ind*_/*H*_*E*_ is independent of read depth, number of individuals sampled, and the allele frequency. However, all of these factors influence the variance of the estimate, and low minor allele frequency especially can bias it upwards (Fig. 5-6). As there is no generalized formula to estimate the variance of a ratio, the variance of *H*_*ind*_/*H*_*E*_ cannot be estimated mathematically. Moreover, sequencing error inflates the estimate at low minor allele frequency (Fig. 5), and polyRAD cannot account for sequence quality scores or alignment quality scores since it only imports allelic read depth. We therefore recommend simulating data for Mendelian loci given the ploidy, inbreeding, sample size, sequencing error rate, and distribution of read depth and allele frequency observed in the dataset of interest. The distribution of *H*_*ind*_/*H*_*E*_ across simulated loci then can be used to determine cutoff values for filtering loci in the empirical dataset. The *ExpectedHindHe* and *ExpecteHindHeMapping* functions are available in polyRAD for this purpose, and suggest cutoffs for filtering loci in order to retain 95% of Mendelian loci. Depending on the downstream application, we recommend considering the number of markers needed versus the importance of marker quality when determining thresholds for read depth, allele frequency, and *H*_*ind*_/*H*_*E*_.

*H*_*ind*_/*H*_*E*_ is more useful for detecting paralogs when haplotypes are treated as alleles (i.e. loci can be multiallelic), as opposed to when all loci are treated as biallelic SNPs, simply due to the fact that multiallelic markers are more information-rich than biallelic markers for the same distribution of minor allele frequencies. We observed that, for the same set of SNPs in *M. sacchariflorus*, the median value of *H*_*ind*_/*H*_*E*_ per locus was very similar regardless of whether they were phased into haplotypes within the span of a single RAD tag, but the variance in *H*_*ind*_/*H*_*E*_ was about 20% higher for SNPs vs. haplotypes (Additional File 1: Fig. S3). This improved power and information content is why polyRAD generally imports multiallelic, haplotype-based genotypes rather than SNPs as the default. Other methods for marker calling in highly duplicated genomes have also benefitted from the use of haplotype information [11, 23], and multiallelic markers have been found to be advantageous over biallelic SNPs for linkage mapping in polyploids [24]. It should be noted that in this study we only phased SNPs that were certain to have originated from the same sequencing reads based on physical linkage and read depth. The *H*_*ind*_/*H*_*E*_ statistic cannot be estimated using haplotypes spanning longer distances, given that read depth will vary from locus to locus within haplotype.

### Uses of the *H*_*ind*_/*H*_*E*_ statistic

We anticipate locus-filtering to be the most common application of the *H*_*ind*_/*H*_*E*_ statistic, with major advantages being that it is not biased by read depth or allele frequency and can be estimated prior to genotype calling. We demonstrate that it is similar to *H*_*O*_/*H*_*E*_ in effectiveness for filtering paralogs, with substantial savings on computational time (Table 2). We should note that our *H*_*O*_/*H*_*E*_ estimates used Bayesian genotype calls from polyRAD, which mitigate the underestimation of observed heterozygosity as compared to naïve genotype calls [12]. Stringency of filtering should depend on the genotype quality needed for downstream analysis; for example, parentage analysis and QTL mapping are sensitive to genotyping errors, whereas genome-wide association studies and estimations of population structure from principal components analysis are less sensitive. Missing data rate, median read depth, and minor allele frequency are common criteria that should be used in combination with *H*_*ind*_/*H*_*E*_ to determine which loci to retain for downstream analysis. In our empirical dataset, we found the loci ranging in depth from five to 100 had the best distribution of *H*_*ind*_/*H*_*E*_ (Fig. 1), but a higher minimum depth may be required for applications that require accurate genotype calling, and the optimal maximum depth used in filtering depends on the overall depth of the dataset. The use of observed heterozygosity, read depth ratios within genotypes, and number of haplotypes per individual are redundant with *H*_*ind*_/*H*_*E*_ and unnecessary if it has already been used for filtering. In addition to its use for detecting paralogs in highly duplicated genomes, *H*_*ind*_/*H*_*E*_ can be used for marker filtering in less duplicated genomes where occasional paralogs are still an issue.

Additionally, in any species, markers with low values of *H*_*ind*_/*H*_*E*_ (e.g. below the 95% confidence interval generated by simulated data) are likely to have null alleles, high overdispersion, or other technical issues and should generally be removed from the dataset. We found that using *H*_*ind*_/*H*_*E*_ to filter our *M. sacchariflorus* dataset impacted minor allele frequency and proportion of markers in genes in ways consistent with the removal of markers with null alleles, collapsed paralogs, or false alleles due to sequencing error (Table 2 and Fig. 3).

Although less accurate for determining ploidy than techniques such as flow cytometry, when averaged within individuals, *H*_*ind*_/*H*_*E*_ can be used to identify individuals whose ploidy might deviate from expectations and should be confirmed. If flow cytometry is not an option, several other tools exist for the estimation of ploidy directly from next-generation sequencing data [25]. Lastly, *H*_*ind*_/*H*_*E*_ could be potentially useful for improving reference genome assemblies, increasing the value of complementing a de novo assembly with a resequencing or genotyping-by-sequencing effort in a large population or diversity panel. Regions of the reference genome that contain collapsed paralogs are expected to have inflated *H*_*ind*_/*H*_*E*_ values, which could be visualized in a smoothed plot of *H*_*ind*_/*H*_*E*_ vs. alignment position.

At a minor allele frequency of 0.05, a read depth of five or higher is sufficient to estimate *H*_*ind*_/*H*_*E*_ with minimal variance (Fig. 6). It is notable that a read depth of five is too low to call genotypes with confidence, to some extent in diploids but especially in polyploids. However, using the *H*_*ind*_/*H*_*E*_ statistic, such low depth data are useful for a variety of applications such as identification of outlier individuals in terms of ploidy and hybridity, estimation of inbreeding, identification of loci with technical issues, and assessment of reference genome quality. This in turn can enable researchers to reduce sequencing costs by generating preliminary, low-depth datasets to evaluate these issues before (or instead of) sequencing more deeply.

## Conclusions

Here we introduce the *H*_*ind*_/*H*_*E*_ statistic, which can be used for evaluating marker and sample quality in genotyping-by-sequencing datasets for a variety of downstream applications. We demonstrate that reads from paralogous loci cause the statistic to be above the expected value, whereas technical issues such as overdispersion and null alleles cause the statistic to be below the expected value. In typical datasets (hundreds of individuals, read depth above five) the statistic has sufficiently low variance to be useful for filtering loci. The polyRAD R package can estimate *H*_*ind*_/*H*_*E*_, suggest filtering cutoffs based on simulated data, and perform genotyping after filtering.

## Materials and Methods

### Implementation in polyRAD

Functions for estimating *H*_*ind*_/*H*_*E*_ and *H*_*ind*_/*H*_*E*.*map*_ are available in polyRAD v1.2 and later, and are named *HindHe* and *HindHeMapping*, respectively. Both utilize an internal Rcpp function for fast calculation, take a *RADdata* object as input, and return a matrix of values, with samples in rows and loci in columns. The mean value across rows can then be used to get a per-sample estimate, for identifying individuals that are interspecies hybrids or unexpected ploidies. The mean value across columns can be used to get a per-locus estimate for filtering loci. Additionally, polyRAD v1.5 and later includes the *ExpectedHindHe* and *ExpectedHindHeMapping* functions, which simulate data to emulate the sample size, allele frequency distribution or parental genotypes, and read depth distribution of an empirical dataset, and return the distribution of *H*_*ind*_/*H*_*E*_ as if all loci were Mendelian, giving the user reasonable thresholds to use for filtering loci.

PolyRAD v1.6 is currently available on CRAN, and can be installed using *install*.*packages(“polyRAD”)*.

### Datasets for testing

Two types of datasets were used to test *H*_*ind*_/*H*_*E*_: (1) empirical data from a diversity panel of *Miscanthus sacchariflorus*, and (2) simulated datasets of diversity panels and of biparental F1 mapping populations. Previously published RAD-seq data for an *M. sacchariflorus* diversity panel [22] were used for the empirical tests. All species in the *Miscanthus* genus share an ancient genome duplication, increasing the chromosome number to 19 from the base of 10 in the Andropogoneae tribe [26–28]. Moreover, some populations of *M. sacchariflorus* display autotetraploidy in addition to this genome duplication (4x = 76) [22, 29], allowing us to test our algorithm in situations where tetrasomic inheritance is expected, in addition to the more typical disomic inheritance. *Miscanthus* is also highly heterozygous due to being wind-pollinated and self-incompatible [30], thus heterozygosity cannot be used to identify paralogs as easily as it could in an inbred crop species. Together, these factors make *M. sacchariflorus* an ideal test case.

To compare values of *H*_*ind*_/*H*_*E*_ in putatively Mendelian markers versus collapsed paralogs, markers were called from the same dataset using either *Miscanthus sinensis* or *Sorghum bicolor* as a reference because *M. sinensis* has a whole genome duplication with respect to *S. bicolor*. Raw sequence reads from *M. sacchariflorus* were processed by the TASSEL-GBSv2 pipeline [31] to identify unique tag sequences and their depths in all individuals. Tag sequences were then aligned to the *Miscanthus sinensis* v7.1 reference genome [32] and the *Sorghum bicolor* v3.1.1 reference genome [33] using Bowtie 2 [34]. The tag manager feature of TagDigger [35] was used to process the SAM files, recording the alignment location of each tag in both reference genomes. Tag alignment locations within the *S. bicolor* reference were retained for further analysis if they corresponded to two alignment locations in the *M. sinensis* reference matching the known synteny between chromosomes. Under this filtering, 239,501 tags were retained at 18,402 *S. bicolor* alignment locations corresponding to 36,804 *M. sinensis* alignment locations, in a set of 356 diploid and 268 tetraploid individuals. *H*_*ind*_/*H*_*E*_ was then estimated per-locus in polyRAD for both the *M. sinensis* and *S. bicolor* alignments.

To compare the variance of *H*_*ind*_/*H*_*E*_ when biallelic SNPs were used versus multiallelic, haplotype-based markers, the TASSEL-GBSv2 pipeline was used to call SNP variants from *M. sacchariflorus* and export them to VCF. Markers from chromosome 1 were imported to polyRAD using *VCF2RADdata*, with and without the option to phase SNPs into haplotypes, yielding 3710 and 10,458 loci, respectively. The phasing performed by *VCF2RADdata* only phases SNPs that are certain to have originated from the same reads based on allelic read depth and physical distance. *H*_*ind*_/*H*_*E*_ was then estimated by locus in polyRAD separately for diploids and tetraploids.

Simulated diversity panel datasets were generated in order to assess the effect of minor allele frequency, sample size, read depth, sequencing error, overdispersion, inbreeding, ploidy, and null alleles on variance and bias of the *H*_*ind*_/*H*_*E*_ statistic, using the *SimGenotypes* and *SimAlleleDepth* functions in polyRAD v1.6. See Clark et al. [12] (Eqn. 2) for a definition of the overdispersion parameter; lower values result in allelic read depths that deviate further from the ratios expected based on allelic dosage. Three sets of data were simulated. (1) Minor allele frequencies of 0.01, 0.05, and 0.1; sample sizes of 100, 500, and 1000; and genotype read depths of 2, 5, 10, 20, 50, and 100 were simulated in all combinations under diploidy and tetraploidy, with no inbreeding, a sequencing error rate of 0.001, and an overdispersion parameter of 20. For each combination, 5000 biallelic loci were simulated, as well as 5000 collapsed paralogs that each consisted of two Mendelian loci combined. (2) Minor allele frequencies of 0.01 and 0.05, overdispersion spanning all integers from 5 to 20, sequencing error rates of 0 and 0.001, and inbreeding (*F*; the probability that two locus copies in an individual are identical by descent) spanning all intervals of 0.1 from 0 to 1 were simulated in all combinations under diploidy and tetraploidy, with a sample size of 500 and a read depth of 20. For each combination, 20,000 biallelic loci were simulated. (3) Minor non-null allele frequencies of 0.01 and 0.05 and null allele frequencies of 0.01, 0.05, 0.1, and 0.2 were simulated in all combinations under diploidy and tetraploidy, with a sample size of 500, a read depth of 20, a sequencing error rate of 0.001, overdispersion of 20, and no inbreeding. For each combination, 5000 triallelic (with one allele being null, i.e. having all of its reads discarded) loci were simulated.

Simulated F1 mapping population datasets were generated in order to assess the effect of ploidy and marker type on variance of the *H*_*ind*_/*H*_*E*_ statistic. For diploids, testcross (homozygote x heterozygote) and F2 (heterozygote x heterozygote) markers were evaluated. For tetraploids, simplex x nulliplex (AAAB x AAAA), duplex x nulliplex (AABB x AAAA), triplex x nulliplex (ABBB x AAAA), simplex x simplex (AAAB x AAAB), simplex x duplex (AAAB x AABB), simplex x triplex (AAAB x ABBB), and duplex x duplex (AABB x AABB) markers were evaluated. For each marker type, 5000 biallelic markers were simulated in a population with 500 offspring, with a read depth of 20, a sequencing error rate of 0.001, and overdispersion parameter of 20.

To evaluate effectiveness of various approaches for filtering paralogs, 1000 Mendelian loci and 1000 collapsed paralogs were simulated in 200 diploid and 200 tetraploid individuals each at three levels of inbreeding. Number of alleles was evenly distributed from two to eight in Mendelian loci. Allele frequency was sampled from a gamma distribution with shape of 0.3 and scale of 1, divided by 10 and added to 0.01 to ensure a minimum minor allele frequency, given that allele frequency filtering is typically performed during variant calling and/or data import. One allele frequency at each locus was generated as one minus the sum of all other allele frequencies, to emulate the typical situation of one common allele and one or more rare alleles. Genotypes were simulated from the allele frequencies assuming an inbreeding coefficient (*F*) of 0.1, 0.5, or 0.9. Mean read depth per locus was drawn from a gamma distribution with a shape of 3.2 and scale of 8. Read depth at individual genotypes was then drawn from a gamma distribution with the locus depth / 10 as the shape, and a scale of 10. Allelic read depth was simulated assuming an overdispersion parameter of 20 and a sequencing error rate of 0.001. Collapsed paralogs were simulated in the same way, but with number of alleles per locus ranging from one to eight, and two random loci being combined to form a collapsed paralog.

### Comparison with other approaches

To call genotypes for the *H*_*O*_/*H*_*E*_ and Z-score [19] approaches, the *IterateHWE* function in polyRAD was used with default parameters to obtain genotype probabilities, and then *GetProbableGenotypes* was used to get discrete genotypes, with genotypes set to missing if allele copy numbers did not add up to the ploidy. To extend its use to polyploids, *H*_*O*_ was estimated as the probability that two alleles sampled from a genotype without replacement would be different from each other, averaged across individuals within a locus. The Z-score approach [19] was originally only defined for biallelic markers in diploids. To extend it for multiallelic markers and polyploid species, for each marker genotypes with *ploidy – 1* copies of the most common allele (i.e. the heterozygous genotype class expected to be most common) were identified, and allelic read depth summed across those samples. Deviation of read depth of the most common allele from the expected ratio was then estimated as a Z-score:

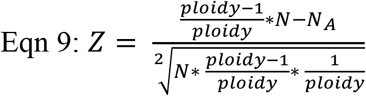

Where *N* is the total read depth across all samples in the given heterozygous genotype class, and *N*_*A*_ is the read depth of the common allele summed across those same samples. The number of haplotypes per genotype was counted as the number of haplotypes with read depth of three or higher, following Willis et al. [17].

## Declarations

### Ethics approval and consent to participate

Not applicable

### Consent for publication

Not applicable

### Availability of data and materials

Raw sequence reads for *M. sacchariflorus* are available on the NCBI Sequence Read Archive under accessions SRP026347, SRP048207, SRP063572, and SRP087645. Genotype calls and read depths for *M. sacchariflorus* are available on the Illinois Data Bank at https://doi.org/10.13012/B2IDB-8170405_V1. Tag alignments and counts for *M. sacchariflorus* using the *M. sinensis* and *S. bicolor* reference genomes are available on the Illinois Data Bank at https://doi.org/10.13012/B2IDB-4814898_V1. All scripts for testing the *H*_*ind*_/*H*_*E*_ statistic are available on GitHub at https://github.com/lvclark/paralog_id, archived on Zenodo at https://doi.org/10.5281/zenodo.5425343.

Project name: polyRAD

Project home page: https://github.com/lvclark/polyRAD

Archived version: https://doi.org/10.5281/zenodo.1143744

Operating system: Platform independent

Programming language: R, C++ via Rcpp

Other requirements: R ≥ 3.5.0; CRAN packages fastmatch, Rccp, and stringi; Bioconductor package pcaMethods

License: GNU GPL (≥2)

Any restrictions to use by non-academics: None

### Authors’ contributions

LVC designed the *H*_*ind*_/*H*_*E*_ statistic, wrote the polyRAD software, performed the analysis, and wrote the manuscript. WM performed literature review and tested the software and statistic. AEL gave statistical advice. EJS provided the *M. sacchariflorus* datasets. All authors read and approved the final manuscript.

## Acknowledgements

We thank Jiale He for testing various methods to detect paralogous loci in *Miscanthus*.

