## Additional File 1 (Supplementary Figures) for "A population-level statistic for assessing Mendelian behavior of genotyping-by-sequencing data from highly duplicated genomes"

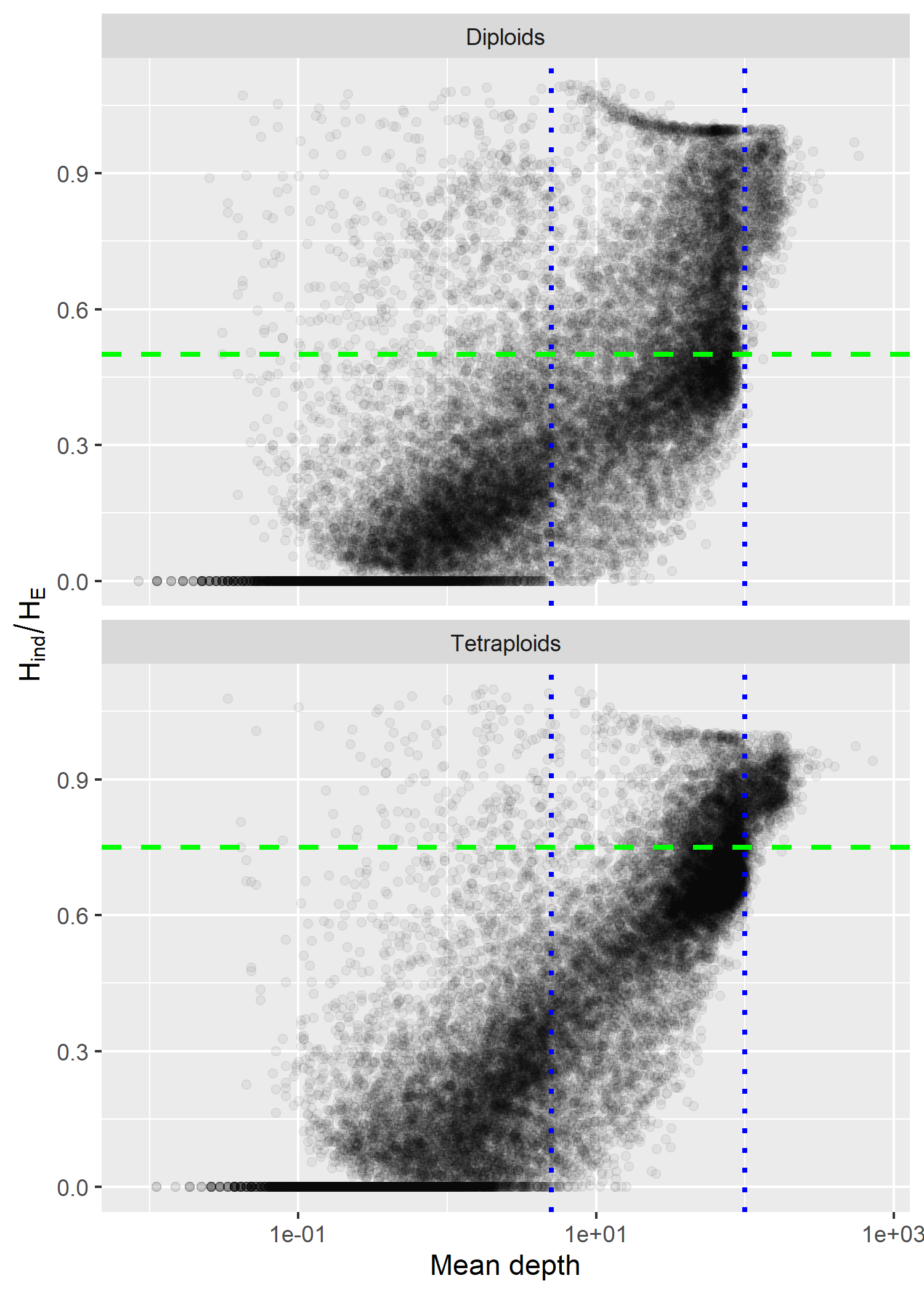

**Figure S1. Relationship between *H_ind_*/*H_E_* statistic and mean sequence read depth per locus.** Loci were called across 356 diploid and 268 tetraploid *Miscanthus sacchariflorus* based on alignments to the *M. sinensis* reference genome. The expected value for a Mendelian locus in Hardy-Weinberg equilibrium is shown with a green dashed line. Cutoffs of 5 and 100 used for Fig. 1 in the main manuscript are shown with blue dotted lines.

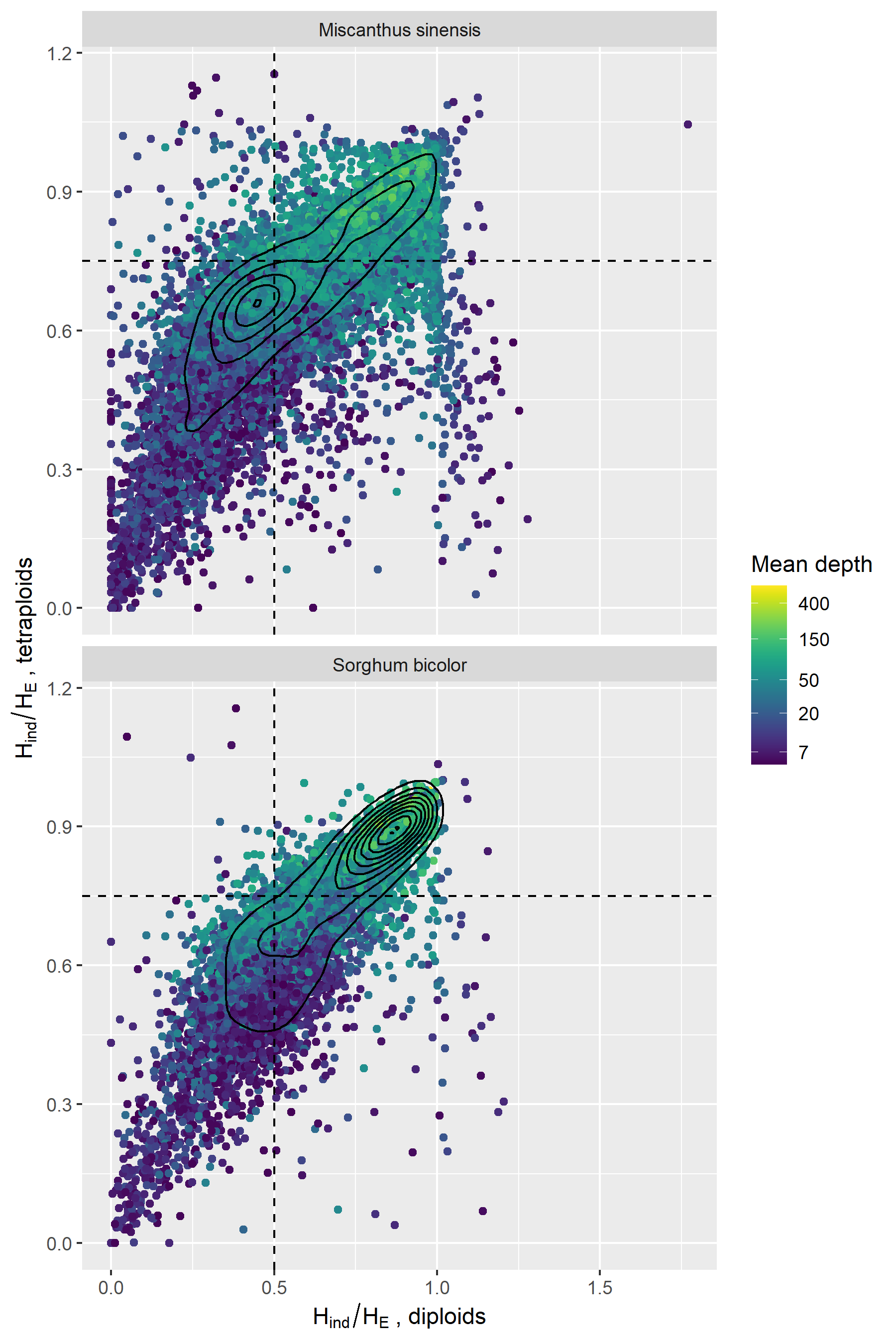

**Figure S2. Correlation of *H_ind_*/*H_E_* between diploid and tetraploid datasets of *Miscanthus sacchariflorus*.** Loci with a mean read depth below five were omitted, leaving 11,516 loci aligned to the *M. sinensis* reference and 8,820 loci aligned to the *Sorghum bicolor* reference. Expected values for Mendelian loci under Hardy-Weinberg equilibrium are shown with dashed lines. Solid lines are 2D density plots.

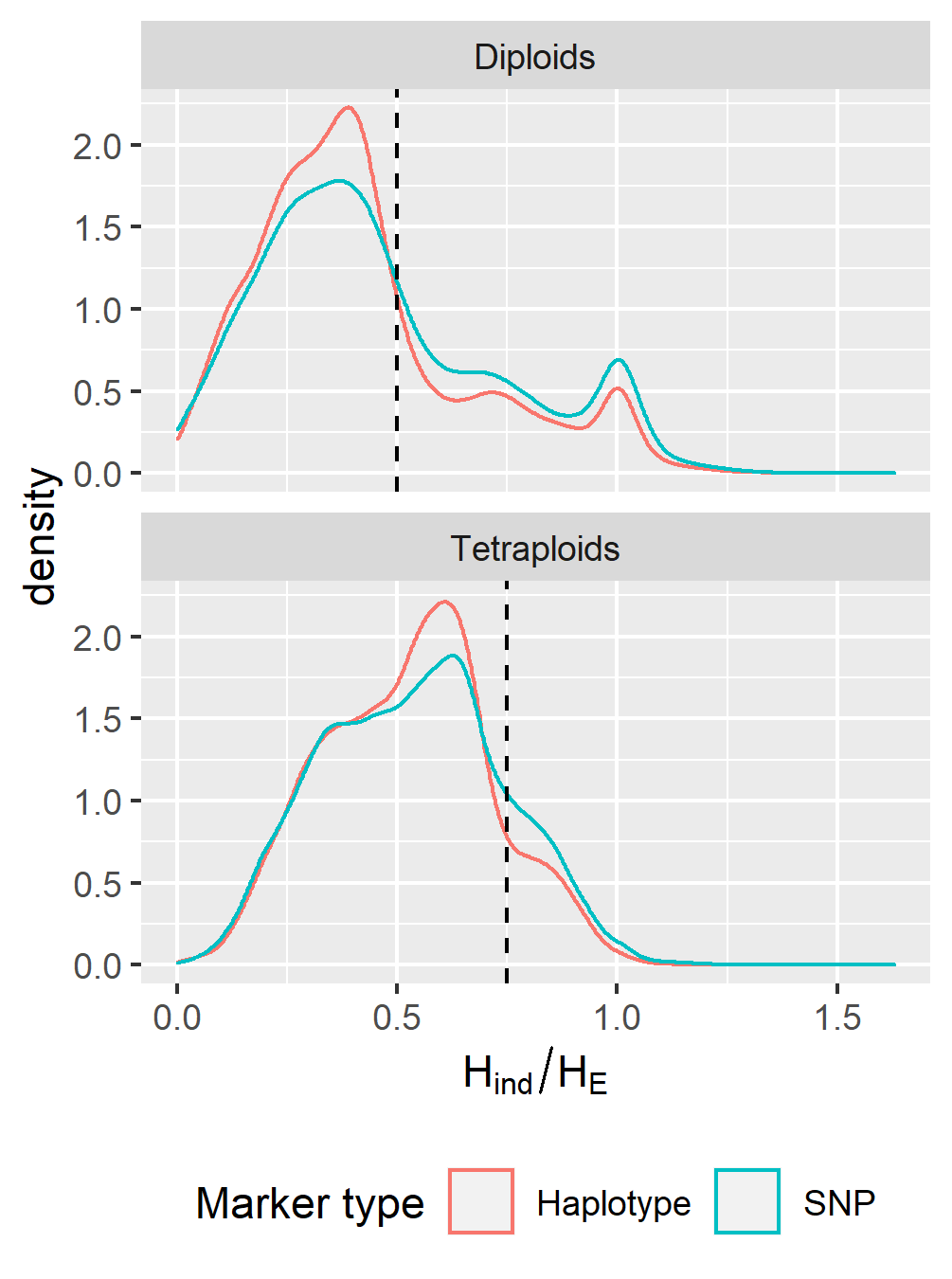

**Figure S3. Distribution of *H_ind_*/*H_E_* among SNPs and haplotype-based loci in diploid and tetraploid *M. sacchariflorus*.** The expected value for Mendelian loci under Hardy-Weinberg equilibrium is shown with the dashed line. The statistic was estimated for 10,458 SNPs, as well as those same SNPs phased into 3710 haplotype-based, multiallelic loci.

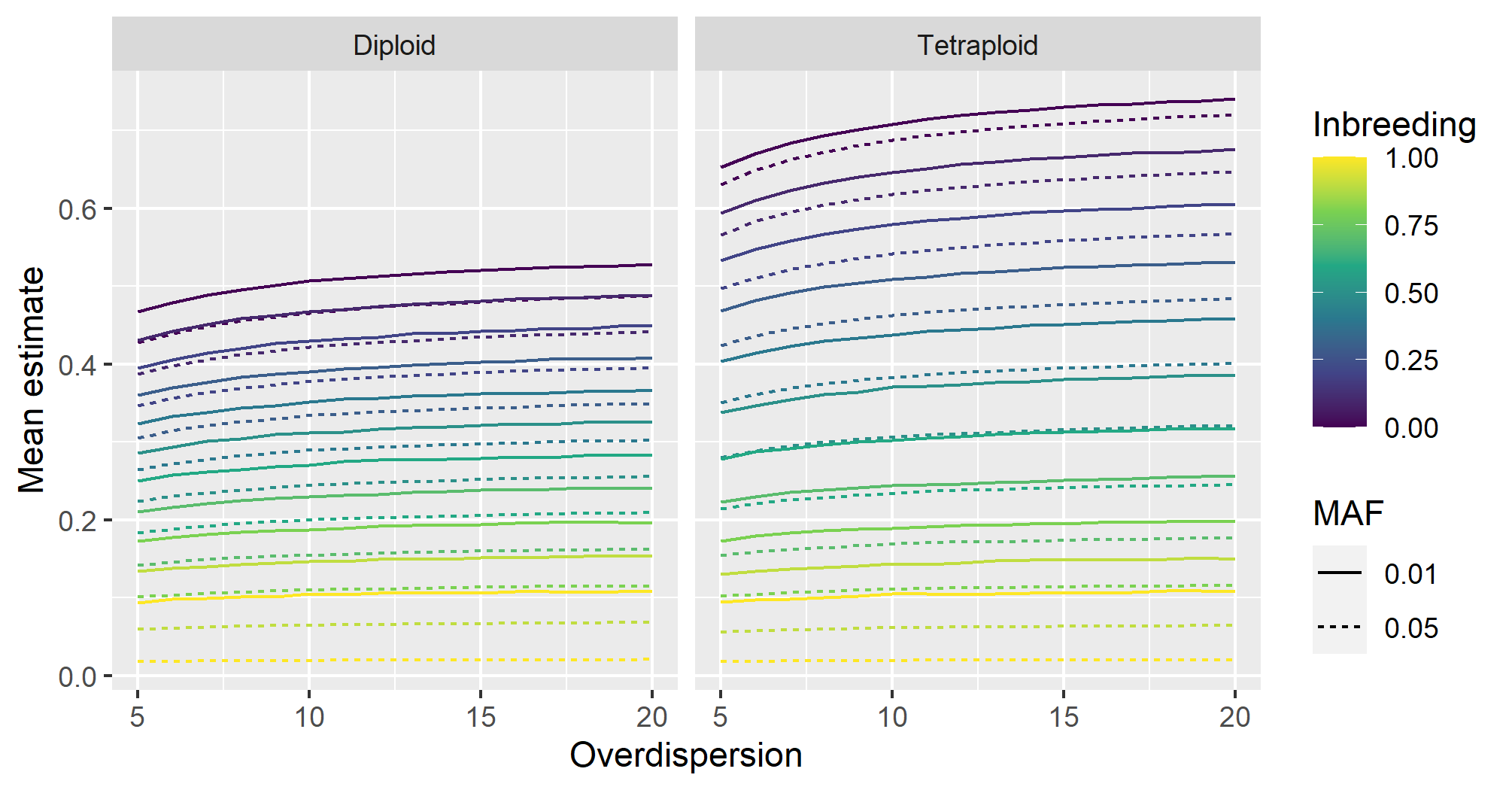

**Figure S4. Mean estimates of *H_ind_*/*H_E_* in simulated datasets under varying inbreeding and overdispersion.** For each ploidy, inbreeding level, overdispersion level, and minor allele frequency (MAF), 20,000 biallelic loci with a read depth of 20 and a sequencing error rate of 0.001 were simulated across 500 individuals, and allelic read depths were used for estimation of *H_ind_*/*H_E_*.

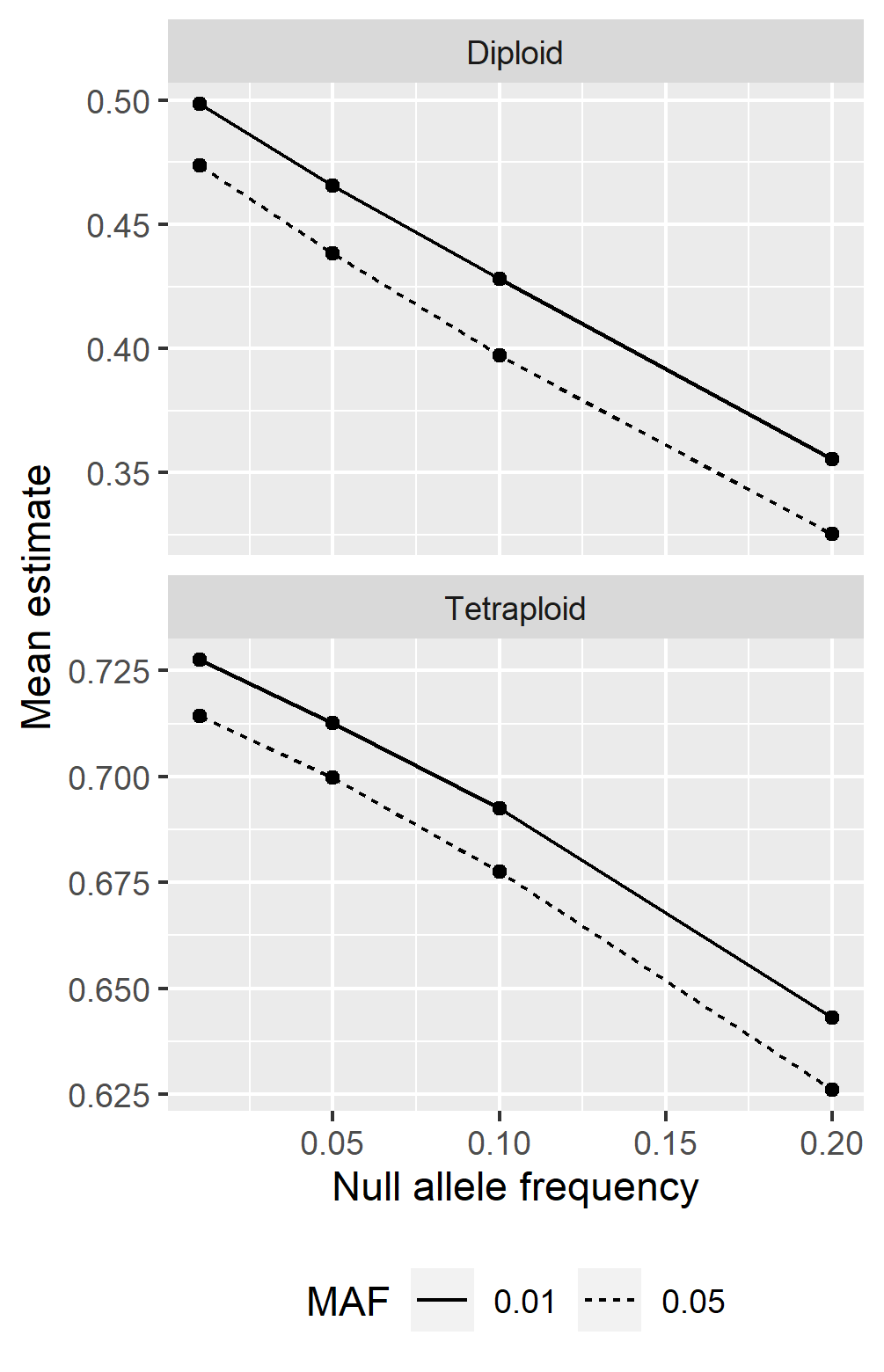

**Figure S5. Mean estimates of *H_ind_*/*H_E_* under varying null allele frequencies.** For each ploidy, null allele frequency, and non-null minor allele frequency (MAF), 5000 loci with a read depth of 20, overdispersion of 20, and sequence error rate of 0.001 were simulated across 500 individuals in a diversity panel or natural population.
